# Non-aging despite high mutation rate – genomic insights into the evolution of *Hydra*

**DOI:** 10.1101/2022.05.18.492491

**Authors:** Arne Sahm, Konstantin Riege, Marco Groth, Martin Bens, Johann Kraus, Martin Fischer, Hans Kestler, Christoph Englert, Ralf Schaible, Matthias Platzer, Steve Hoffmann

**Author notes:** shared senior-authorship (Matthias Platzer, Steve Hoffmann).

## Abstract

*Hydra* is a genus of freshwater polyps with remarkable regeneration abilities and a non-senescent phenotype under laboratory conditions. Thus, this animal is particularly interesting for aging research. Here, we gained insights into *Hydra’s* recent genetic evolution by genome sequencing of single cells and whole individuals. Despite its extreme longevity, *Hydra* does not show a lower somatic mutation rate than humans or mice. While we identify biological processes that have evolved under positive selection in animals kept in optimal laboratory conditions for decades, we found no signs of strong negative selection during this tiny evolutionary window. Interestingly, we observe the opposite pattern for the preceding evolution in the wild over a longer time period. Moreover, we found evidence that *Hydra* evolution in captivity was accompanied and potentially accelerated by loss of heterozygosity. Processes under positive selection in captive animals include pathways associated with *Hydra*’s simple nervous system, its nucleic acid metabolic process, cell migration, and hydrolase activity. Genes associated with organ regeneration, regulation of mRNA splicing, histone ubiquitination, and mitochondrial fusion were identified as highly conserved in the wild. Remarkably, several of the processes under strongest selection are closely related to those considered essential for the exapted, i. e. not brought about by natural selection, feature: *Hydra’s* non-aging.

## Introduction

*Hydra* is a genus of small freshwater polyps with a predicted laboratory lifespan of 1.400 years for *Hydra magnipapillata* (Jones, et al. 2014). Thus, it is often referred as “non-senescent” (Schaible, et al. 2015; Danko, et al. 2015; Klimovich, et al. 2018) “non-aging” (Sturm, et al. 2017; Holtze, et al. 2021) or “immortal” (Müller 1996; Boehm, et al. 2012; Domazet-Lošo, et al. 2014). This controversially discussed attribution is based on the observation that, in stark contrast to most other animals, all *Hydra* species escape senescence under laboratory conditions and while reproducing asexually (Schaible, et al. 2014). This extraordinary longevity seems to be linked to a high regenerative capacity. The animals can replace any part of their body when dissected into multiple fragments. Each part usually regrows into a new organism within two to three days (Fujisawa 2003; Tomczyk, et al. 2015) as shown for the most extreme case of only 1% of the body mass (Shimizu, et al. 1993). These abilities seem to be associated with the simplicity of *Hydra’s* body plan and the fact that most cells are continually proliferating stem cells (Holstein, et al. 1990). Two main stem cell lineages have been described in *Hydra:* faster cycling interstitial cells (I-cells) and slower dividing epithelial cells (E-cells). The latter can be further subdivided into ectodermal and endodermal cells. While I-cells are multipotent and differentiate to neurons, nematocytes, gland cells and gametes, E-cells are unipotent forming the body column and the extremities. Combined the two lineages can differentiate into all *Hydra* cell types. They renew all cells of an individual by continuous replacement within three weeks (Bosch 2009; Hemmrich, et al. 2012; Domazet-Lošo, et al. 2014; Siebert, et al. 2019). Closely linked to the constant regrowth is *Hydra’s* ability of asexual reproduction by budding. During this process, a bud containing one fifth of the approximately 50,000-100,000 cells of an adult individual is formed and detached (Dańko, et al. 2015; Tomczyk, et al. 2015). *Hydra* reproduces asexually in warm water under nutrient-rich conditions, which is fulfilled when kept in captivity. Under less favorable conditions, however, *Hydra* regularly switches to sexual reproduction (Martínez and Bridge 2012; Kaliszewicz and Lipińska 2013). Soon after fertilization *Hydra* embryonal cells differentiate into E- and I-cells that will behave and evolve as separate cell lineages within the clonal pedigree of budded individuals.

To shed light on these phenomena from a molecular perspective, an increasing number of fundamental studies and resources have been published in recent years. These include a low-coverage and therefore quasi mono-allelic genome reference (Chapman, et al. 2010) and transcriptome sequences (Wenger and Galliot 2013), inducible methods for gene knockdown (Watanabe, et al. 2014; Vogg, Beccari, et al. 2019), topological analyses of morphogenesis (Maroudas-Sacks, et al. 2021), as well as quantitative gene expression data from whole individuals and single cells (Krishna, et al. 2013; Petersen, et al. 2015; Siebert, et al. 2019).

However, to this date, evolutionary forces affecting mutation rate and selection remained unexplored. Importantly, *Hydra’s* non-senescent phenotype is not likely to be the result of natural selection - so called exaptation (Stephen Jay Gould, Elisabeth S. Vrba: *Exaptation – a missing term in the science of form*. In: *Paleobiology*. Band 8, Nr. 1, 1982, S. 4–15) - towards a longer lifespan but rather the consequence of features that evolved to increase survival under harsh environmental conditions (Schaible et al. 2014). In the wild, due to a high extrinsic mortality, *Hydra’s* average lifespan is counted in weeks rather than in years (Schaible et al. 2015). The non-senescent phenotype is only observable under benign laboratory conditions, i.e., absence of extrinsic mortality risks, steady feeding and optimal temperature. As a by-product of *Hydra’s* regeneration capabilities, a high cell turnover and stem cell omnipotence can be recorded (Schaible et al. 2017). Non-senescence of *Hydra* individuals, however, does not exclude that stem cells are able to ‘wear out’ and senesce over time, similar to individual bacteria (Stewart et al. 2005, Wang et al. 2010). Whether asymmetric cell division occurred in *Hydra* has still to be investigated.

Somatic mutations are nowadays frequently considered as one or even the major factor of the aging processes (Lopez-Otin, et al. 2013; Vijg 2021; Wolf 2021; Yousefzadeh, et al. 2021). Essentially, it is assumed that increasing DNA damage gradually undermines the integrity of individual cells, followed by tissues, organs, and finally the entire organism. This process leads to its progressive decay and aging-associated diseases.

In this study, we systematically examine Hydra’s mutation rate as well as positive and negative selection processes across different evolutionary time scales using single-cell and whole-animal genome sequencing data of *H. magnipapillata* strain 105.

## Results

To determine mutation rate and to identify signs of selection in the female strain 105 of *H. magnipapillata*, we performed (i) whole-individual genome sequencing of a pedigree, consisting of its mother and 8 daughters (buds) generated at different time points between 2006 and 2018, (ii) single-cell whole genome sequencing of three E- and three I-cells obtained from an individual independently raised in our laboratory and (iii) integration of our results with the currently available draft genome assembly (Chapman, et al. 2010, Fig. 1, Supplement Table S1).

**Fig. 1.**
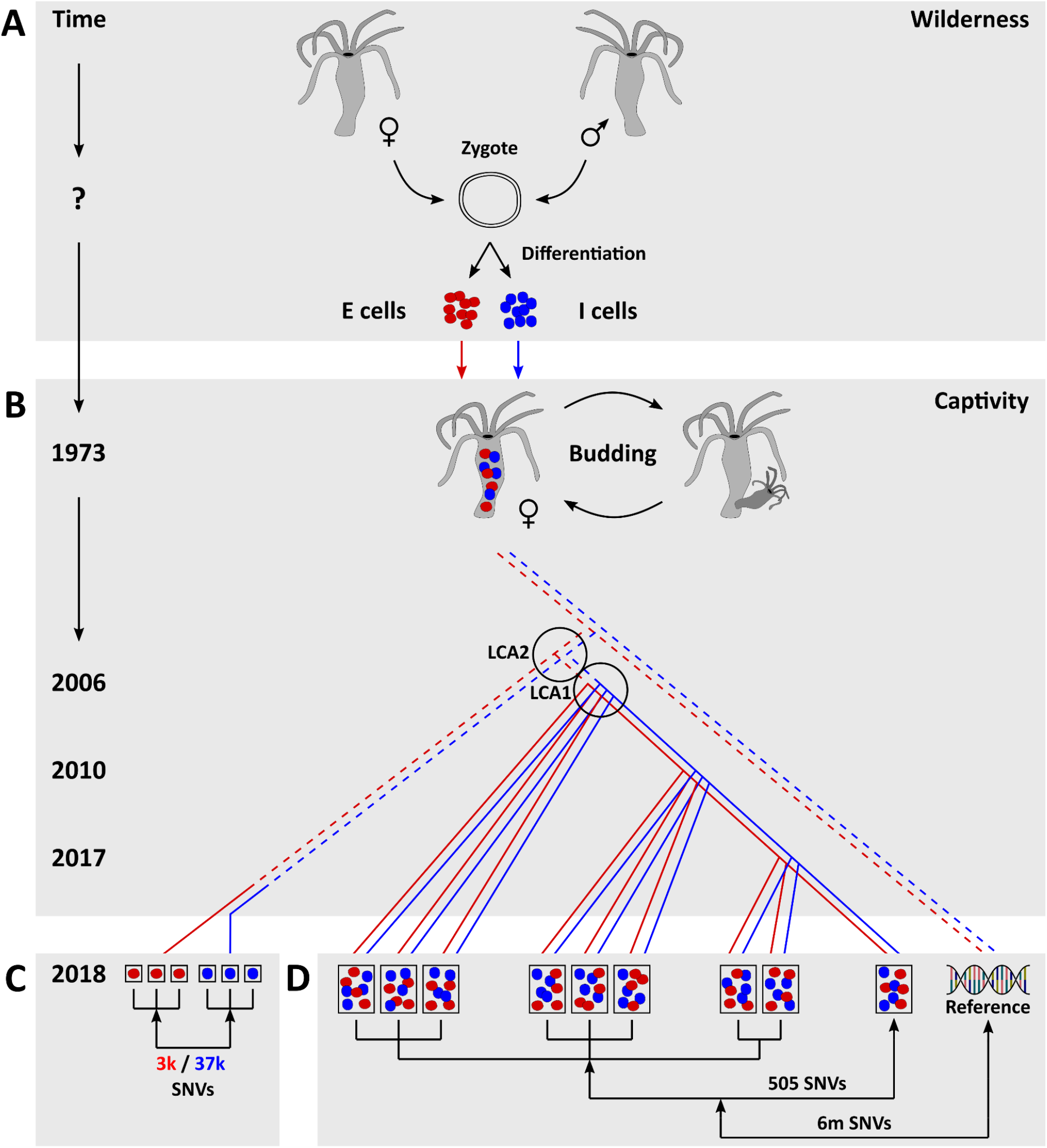
Biological and historical background, experimental setup and pedigree relationships of analyzed data sets of *H. magnipapillata* strain 105. (**A**) The last sexual reproduction, representing the strain’s genetic origin, occurred in the wild. Shortly after fertilization, a *Hydra* zygote differentiates into non-interchangeable epithelial (E-cells, red) and interstitial (I-cells, blue) stem cell lineages. (**B**) The *H. magnipapillata* strain 105 originated from a single female polyp collected by Dr. Tsutomu Sugiyama in September,1973 from a swamp adjacent to the National Institute of Genetics in Mishima, Japan (T. Fujisawa, unpublished information). The strain has since reproduced exclusively asexually by budding and has been propagated in labs worldwide. For generating a reference genome sequence, the strain was re-cloned from a single polyp in the laboratory of Dr. Hans Bode at UC Irvine in 2004 (Chapman, et al. 2010). We obtained the strain from UC Irvine in 2006 and established a pedigree, consisting of its mother and 8 daughters (buds) generated at three time points: 2006, 2010 and 2017. (**C**) In 2018, we used single-cell whole-genome sequencing of three E-cells and three I-cells to identify single nucleotide variants (SNVs) that have accumulated in each of these cells since their common origin (zygote). (**D**) In addition, we performed whole-animal sequencing of nine Hydra individuals, comprising the pedigree mentioned above. We identified SNVs that arose in each daughter in comparison to the mother. To further identify heterozygous genome positions of the strain, we also compared all nine polyps individually with the reference genome sequence.. The total numbers of uniquely identified SNV positions are displayed: in (C) for the comparisons of each E-cell *vs* all I-cells combined (red) and each I-cell *vs* E-cells combined (blue); in (D) for the comparisons of buds *vs* mother and individuals *vs* reference. In our experimental setting, the last common ancestor (LCA) 1 of the 9-individuals-pedigree was the mother at its stage of somatic evolution in 2006. The LCA 2 of our 9-individuals-pedigree and the individual used for single-cell sequencing was budded also in 2006. The last common ancestor of the individuals sequenced by us and those from which the reference genome was derived can be dated back only to a time window of 1974-2004. Pedigree lines: continuous - life history of a Hydra individual; dashed - descant relation including multiple *Hydra* individuals derived from each other by budding.

### Single nucleotide variants are detectable over different short evolutionary time scales

Analyzing our genomic sequencing data (Supplement Text S1), we identified single nucleotide variants (SNVs in three comparisons covering different evolutionary time scales:

1. Within the 9-animal pedigree, we compared each daughter (bud) against the mother resulting in an average of 101 SNVs per bud (505 unique variant positions in total across all buds, Supplement Table S2). In this setting, the last common ancestor (LCA) 1 denotes the mother at her stage of somatic evolution in 2006, i.e. 12 years before sampling (Fig. 1). Our analyses suggest that these variants are almost exclusively a product of founder effects and subsequent accumulation over time. However, we found no evidence for an enrichment of the mother’s mutations in the buds (Supplement Text S1, Supplement Fig. S1A). This comparison covered the shortest examined time scale, from the respective budding events to the sampling in 2018.
2. We identified SNVs that accumulated in single cells since the last sexual reproduction by comparing each of them against the three single cells from the respective other stem cell lineage. The last sexual reproduction happened before 1973 since the analyzed strain was derived from a single female animal captured in that year in the wild. The purely female strain has reproduced exclusively asexually ever since (Sugiyama and Fujisawa 1977; Schaible, et al. 2015). We identified an average of 14,459 SNVs per I-cell and 1,083 SNVs per E-cell. In total, we found 37,379 and 3,184 unique positions affected by mutations in the stem cell lineages of I-cells and E-cells, respectively. Consequently, we observe a 10-fold increase of the mutational load in the faster dividing I-cells. In part, however, the marked difference of SNVs between the lineages can be attributed to the sequencing coverage which was higher in I-cells (Supplement Table S3). To ensure comparability with a previous study (Milholland, et al. (2017), we used the same methodology for sequencing and bioinformatics analysis.
3. We examined each of the sequenced individuals of our 9-individuals-pedigree for deviations from the mono-allelic reference genome (Chapman, et al. 2010). The identified SNV alleles were most likely already present in the two haploid gamete genomes combined in the zygote (Supplement Text S1) that gave rise to the individual caught in 1973 from the wild. These variants thus cover by far the longest of the time scales analyzed and provide information about evolutionary pressures on *Hydra* in the wild. We detected approximately 4.5 million SNVs per comparison and 6 million unique SNV positions in total across all samples (Supplement Table S3).

### Hydra’s mutation rate is not lower than that of human and mouse

By further analyzing the single-cell data, we found an SNV on average every 5,770 bases in I-cells and every 53,492 bases in E-cells. Since replication errors are a major source of genomic variation (Busuttil, et al. 2006), the higher mutational load of I-cells may be explained by their approximately three-fold difference in cell division rate: every 24-30 h and every 3-4 days, respectively (David and Campbell 1972; Campbell and David 1974). Based on these cell division frequencies and the estimation that I-cells and E-cells started to evolve separately approximately 44 years before our sampling in 2018 (Fig. 1), we inferred an average mutation rate per cell division of 1.21*10^−8^ for I-cells and 4.06*10^−9^ for E-cells (Fig. 2A). In each of the six cells studied, the mutation rate in the coding sequence is lower than in the rest of the genome (p=0.031, paired Wilcoxon test), although the differences are smaller for E-cells. On average, the estimated mutation rate per cell division in the coding sequence is 1.02*10^−8^ for I-cells and 3.46*10^−9^ for E-cells (Fig. 2A, Supplement Table S2).

**Fig. 2.**
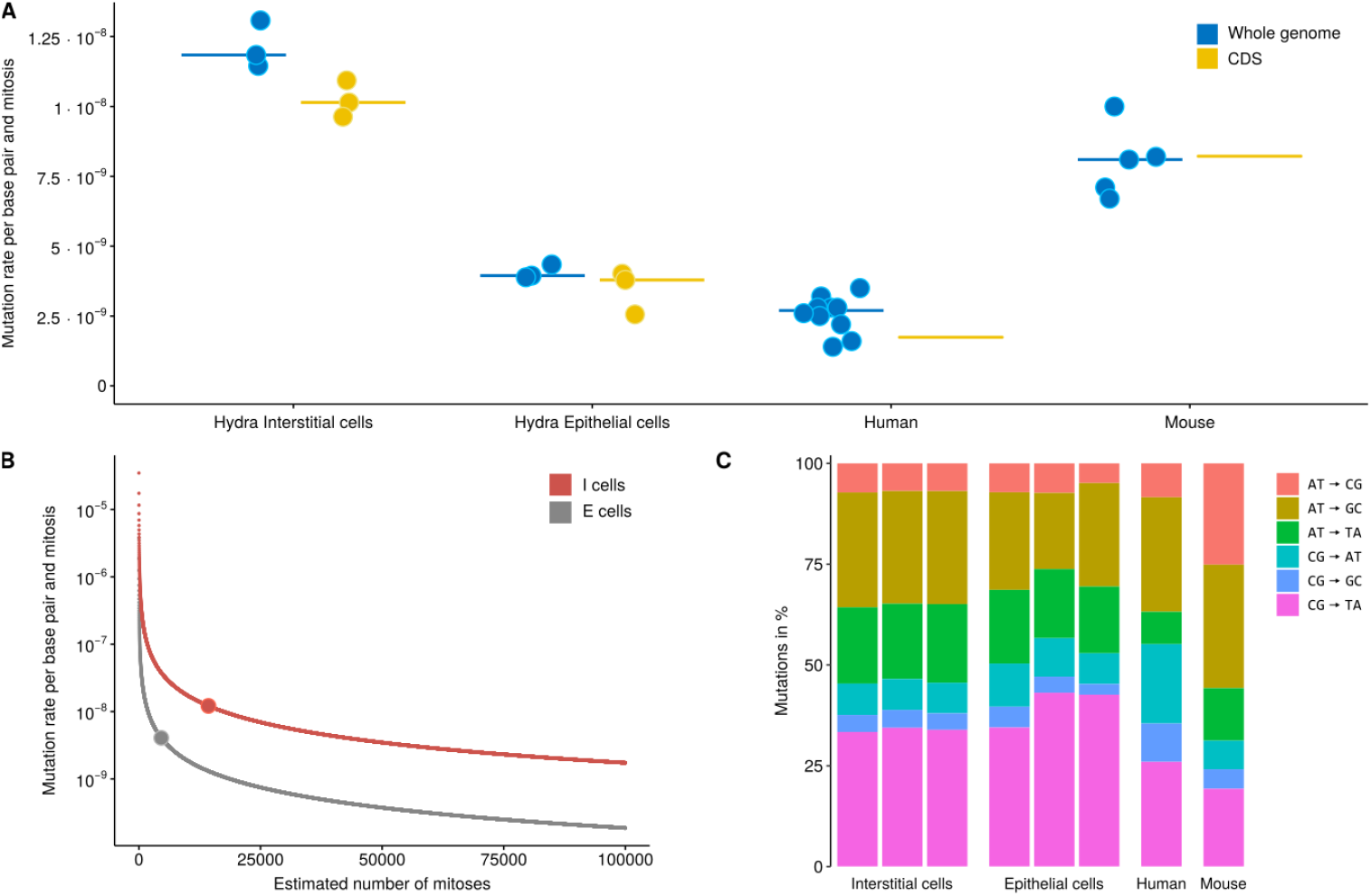
Single-cell whole-genome analysis to determine the mutation rate per cell division in *Hydra*. **(A)** Mutation rates per cell division in *Hydra* I- and E-cells in genome and coding sequences (CDS) compared with literature data for human and mouse dermal fibroblasts (Milholland, et al. 2017). Points represent single sample estimates, lines denote medians (Supplement Table S2; for CDS of human and mouse only averages were published). **(B)** Dependence of mutation rate estimates on the number of cell divisions since the zygote. Dots represent predicted mutation rates given in (A) at the, as estimated by us, numbers of cell divisions that occured within 44 years, i.e. 14,292 and 4,605 divisions for I-cells and E-cells, respectively. **(C)** Mutation spectra in *Hydra* of the three I- and three E-cells compared to literature data for human and mouse dermal fibroblasts (Milholland, et al. 2017).

Despite *Hydra’s* extreme longevity, the estimated mutation rates are not lower than the corresponding literature values for human and mouse (Milholland, et al. 2017) (Fig. 2A). Naturally, the number of cell divisions that occurred since sexual reproduction (14,292 and 4,605 for I-cells and E-cells, respectively, Fig. 2A) is unknown and can only be estimated by a lower bound. Therefore, the mutation rate is likely to be underestimated. However, the influence of the number of cell divisions on the mutation rate estimate’s order of magnitude is clearly limited (Fig. 2B).

Another potential source affecting the prediction of *Hydra’s* mutation rate are spurious mutations caused, e.g., by elevated temperatures during the cell lysis step. Typical for this artifact is a mutation spectrum that is largely dominated by CG→TA transitions due to the possibility of cytosine deamination under high temperatures (Dong, et al. 2017). However, when we compare the mutation spectra in *Hydra* single cells with that of human and mouse (Milholland, et al. 2017) we only find a slightly higher fraction of CG→TA mutations in our data (Fig. 2C). While it cannot be completely ruled out that a fraction of the identified variants might be caused by methodological problems, a great impact of this on overestimating the mutation rate of *Hydra* seems unlikely.

### Multiple appearance and homozygosity of novel SNV alleles indicate recent positive selection

Seeking signs of selection, we investigated how many variants identified in the single cells were found in the nine individuals sequenced as a whole. Of note, in single cells, we are likely to detect almost all SNVs for those positions that pass a certain read coverage threshold (in our case 20). When sequencing whole individuals, however, such coverage in most cases allows detection of an SNV only if it occurs in a significant proportion of the animal’s cells. Therefore, it is interesting that a majority of variants identified in single cells (2nd comparison) were found also in the individuals (3rd comparison). We found a bipartite pattern with most single-cell SNVs being either present in all nine examined individuals or in none (Fig 3a). Given the relationship structure, the mutational events related to those SNVs that occured in all nine individuals must have happened somewhere between the zygote and LCA 2 (Fig. 1). Those single-cell SNVs that cannot be detected at the level of entire individuals, if not representing amplification artifacts, likely occurred late on the branch from LCA 2 to the donor of the single cells. SNVs detected in one to eight individuals may have occurred before or after LCA 2, i.e. due to limitations of our sequencing depth in fact may also be present in more if not all examined individuals.

**Fig. 3.**
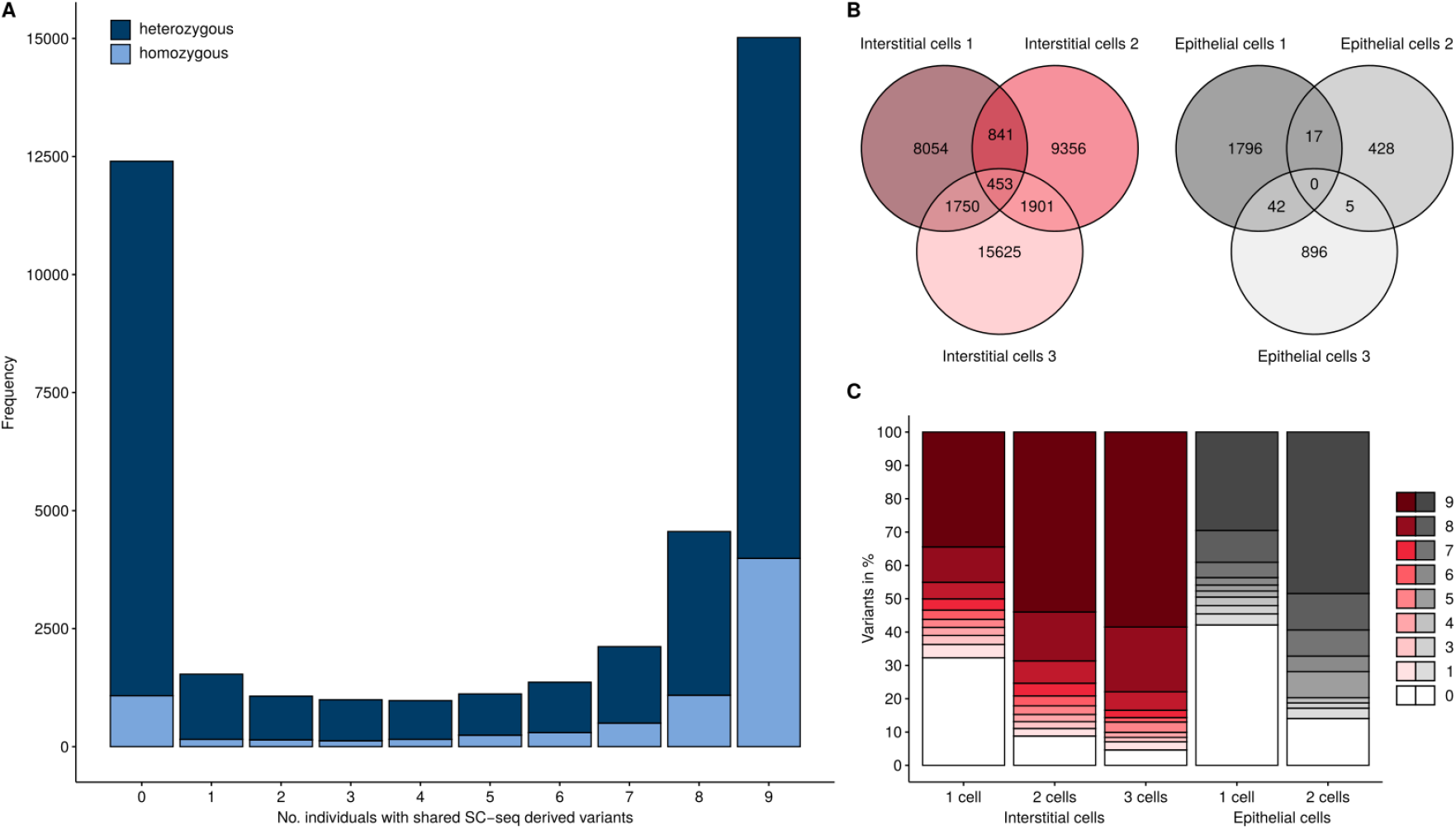
Indicators of positive selection among *Hydra* single-cell SNVs. A) Distribution of hetero- and homozygous single-cell SNVs by their prevalence in the nine individuals. B) Venn analysis of single-cell SNVs in I- and E-cells. No overlap was detected between I-and E-cells. C) Prevalence of single-cell SNVs in 1-9 individuals examined.

By construction, variants found at the single-cell level distinguish both stem cell lineages from each other. Therefore, SNVs found in only one lineage were likely absent in the zygote (Fig.1). If such a variant is found in a significant proportion of the cell population of an entire individual, the respective cell clone might have expanded by somatic positive selection. Given that the three examined I-cells share more variants than the E-cells, positive selection seems to have affected the faster dividing I-cells more strongly (Fig. 3B). In line with that, compared to E-cells, we found I-cell SNVs to be detectable more often at the level of entire *Hydra* individuals (Fig. 3C, p<2.2*10^−16^, Wilcoxon-test). In addition, SNVs that were identified in two I-cells were found in more individuals than those found only in one I-cell (p<2.2*10^−16^). The same relation applies for SNVs found in all three vs two I-cells (p=1.4*10^−3^, Fig 3C). Another indicator of potential events of positive selection is the high percentage of homozygous variants in the single-cell experiment (19%, Fig. 3A). Given the background of lacking regular sister chromosome pairing and recombination during somatic evolution of I- and E-cells, it has to be considered extremely unlikely that by chance so many novel SNV alleles become fixed on both sister chromosomes. If homozygosity of those novel SNV alleles, however, provides considerable fitness advantages to the carrying cells, selection may be considered as the driving force and we could expect those SNV alleles to be particularly prevalent at the level of whole individuals as well.

In line with this, when examining the prevalence of SNVs detected at the single-cell level (2nd comparison) in individuals (3rd comparison), we found homozygous variants exhibiting higher minor allele frequencies than heterozygous ones (median: 0.43 vs 0.28, p<2.2*10^−16^, Wilcoxon-test). Furthermore, the percentage of homozygosity was considerably higher among single-cell SNVs found in all nine individuals compared to those in none of the individuals (27% vs 9%, p<2.2.*10^—16^, Fisher-test).

### Opposite selection patterns manifest before and after a recent drastic environmental change

By genome-wide d_N_/d_S_ analyses across our three examined evolutionary time scales (Fig. 1), we asked how strongly the strain was affected by positive and negative selection. Briefly, the d_N_/d_S_ compares within coding sequences the rate of nonsynonymous variants against that of synonymous SNVs. A d_N_/d_S_ ratio >1 indicates positive selection, of ∼1 neutral evolution, and <1 negative selection (Sahm, et al. 2017). Usually, coding sequences are dominated by negative selection, while events of positive selection are considered rare, episodic exceptions (Yang 2005). In line with this, our single-cell comparison covering the time from the zygote on, i. e. mainly the period in captivity (1973-2018, Fig. 1), showed a genome-wide average d_N_/d_S_ ratio of 0.63. On the longest evolutionary time scale investigated here and mainly reflecting variants accumulated in the wild before the sexual reproduction (3rd comparison of whole-animal sequence data versus reference), negative selection was even more pronounced, with a d_N_/d_S_ ratio of 0.49. The variants, however, that distinguish the examined buds from their mother and accumulated in the shortest examined time scale (2006-18, 1st comparison of whole-animal data of buds vs. mother, Fig. 1), show rather a weak sign of positive selection (d_N_/d_S_ = 1.06).

### Many biological processes under selection are related to Hydra’s regenerative capacity

Finally, we asked which pathways and biological processes were adapted (positive selection, d_N_/d_S_ >1) or strongly conserved (negative selection, d_N_/d_S_ <0.1) during the examined evolutionary times scales. Naturally, this analysis was limited to variants associated with coding sequences of ontologically annotated *Hydra* genes. Of the 505 SNVs identified on the shortest examined time scale (1st comparison of bud data vs. mother) only 15 genes fulfilled these requirements. Therefore, further analysis had to be restricted to the intermediate examined time scale covering mainly the complete time in captivity (1973-2017, 2nd comparison, single-cell data) and the longest examined time scale (3rd comparison, whole-animal data vs reference) including the time before last sexual reproduction in the wild (Fig.1).

Regarding the time in captivity, 28 biological processes show signs of positive selection (Fig. 4A, Supplement Table S4), whereas one process was found to be strongly conserved: positive regulation of multicellular organismal process (GO:0051240, d_N_/d_S_=0.09). Interestingly, for the time in the wild, exactly the opposite pattern was observed: 274 processes display strong negative selection (Fig. 4A, Supplement Table 5), while only one process, i.e. “heterotrimeric G-protein complex” (GO:0005834, d_N_/d_S_=1.30), was found to be adapted. In addition, two processes exhibited an enrichment for nonsense-mutations for the time in the wild (Fisher-test using an FDR threshold of 0.05): “nucleic acid metabolic process” (GO:0090304, FDR=4.5*10^−7^) and “hydrolase activity” (GO:0016787, FDR=7.0*10^−5^). Since both processes were also identified as positively selected for the time in captivity, they may represent evolutionary hot spots in the *Hydra* genome that are subject to continuous adaptation.

**Fig. 4.**
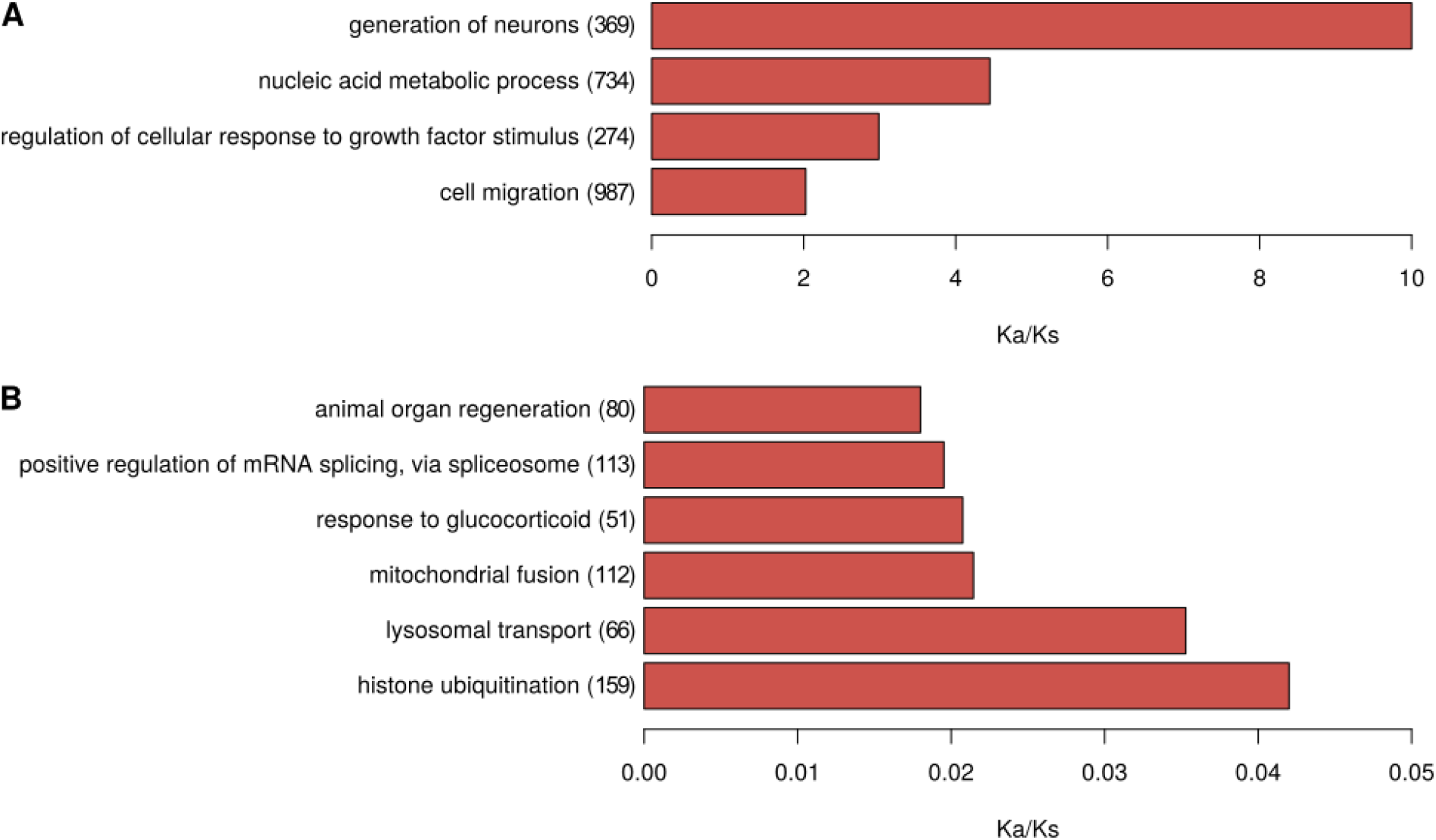
Summary of processes affected (**A**) by positive selection during 44y in captivity after last sexual reproduction and (**B**) by strong negative selection over a longer time scale before last sexual reproduction in the wild. The analysis of single-cell data (**A**) and the comparison of whole-animal data against the *Hydra* reference genome (**B**) resulted in 28 and 274 biological processes affected by positive and strong negative selection, respectively (Supplement tables S4, S5). For this overview, processes were summarized by REVIGO (Supek, et al. 2011). The values in brackets give the number of unique genes that were summarized under the respective term. Strong negative selection was defined by d_N_/d_S_ < 0.1.

## Discussion

Somatic mutations are considered as a major cause of aging (Lopez-Otin, et al. 2013; Vijg 2021; Wolf 2021; Yousefzadeh, et al. 2021). According to the “somatic mutation theory of aging” (Morley 1995), the resulting genetic perturbations can, on the one hand, gradually undermine the integrity of individual cells, tissues and organs. On the other hand, the accumulation of mutations may trigger tumor suppression mechanisms, which, in turn, lead to cell senescence and apoptosis. Eventually, this chain of events erodes the organism’s functions and its ability to regenerate.

The fact that a nonsenescent or only extremely slowly aging organism like *Hydra* does not exhibit a different somatic mutation rate than significantly faster aging species is an argument against the generalization of this theory. However, *Hydra* may be seen as an exception among multicellular organisms since it somewhat resembles a collection of unicellular organisms (Bosch, et al. 2010). Thus, insights into *Hydra* biology may not be directly transferable to most other animals, including mammals.

A potential source of error for the mutation rate is the estimation of cell divisions based on the unknown exact time point of the last sexual reproduction. However, even if we were to assume a much higher number of cell divisions, the mutation rate in Hydra would still not be much lower than in other mammals (Fig 2B). For example, the number of divisions of E-cells would have to be 50% higher than assumed to even approach the mutation rate of humans. Of note, this would imply that the last sexual reproduction would have to have occurred more than 20 years before the capture of the polyp that gave rise to the strain studied here. In the case of I-cells, the number of mitoses would need to be four times higher to even approach the human mutation rate. Consequently, the last sexual reproduction would have occurred at the beginning of the 19^th^ century. Since *Hydra* is known to usually switch to sexual reproduction when conditions are not within a narrow window of optimality, e.g., water temperatures around 19 °C, this seems highly unlikely in a naturallyfluctuating environment (Martínez and Bridge 2012; Kaliszewicz and Lipińska 2013).

Comparing SNV data obtained for the mother and its eight daughters, we found no evidence for the assumption (discussed by Schaible, et al. 2015) that in order to prevent accumulation of damage in the mother, mutated *Hydra* cells may be transmitted via budding to subsequent generations (Supplement Text S1, Supplement Fig. S1A). Instead our data suggest that the constant production of new cells and the continuous removal of cells by differentiation and programmed cell death may be sufficient to prevent accumulation of cells carrying detrimental SNV alleles in individuals at maturity. Moreover, in order to not get lost, novel alleles have to spread out in the *Hydra* cell pool by confining at least a minimal increase in fitness. Consistently, we found in our comparison of whole-animal bud data *vs*. mother a d_N_/d_S_=1.06, i.e. slightly skewed towards positive selection. A more significant fitness increase by novel SNV alleles is rather unlikely, in particular if considering *Hydra* as a “bag” of highly proliferation optimized stem cells - unless environmental conditions change and previously only ‘slightly advantageous’ or even ‘deleterious’ alleles increase cell fitness strikingly. Consistently, we found reliable signs of positive selection among our data only for the intermediate time frame 1973-2018 starting at a drastic change of the environment from wild to laboratory.

Another interesting aspect is the contrasting pattern found regarding the extent of positive and negative selection in the recent evolution of the examined *Hydra* strain with respect to the time in the wild and in captivity. In the wild, negative selection was found in all pathways examined except one; 274 processes were even found to be highly conserved (d_N_/d_S_ <0.1). This part of the results was to be expected, since negative selection dominates in coding sequences - even more so if species are highly adapted to a rather constant environment (Yang 2005). For these reasons it is not surprising that we found extremely little amino acid changes in processes that can be linked directly to highly developed traits of *Hydra*. For example, the regenerative capacity of Hydra is among the most advanced in the animal kingdom (Bosch 2007; Vogg, Galliot, et al. 2019). Therefore, changes in the corresponding genes are likely to decrease fitness. Similarly, the strong purifying selection of positive regulation of multicellular organismal process (GO:0051240) both in the wild and under laboratory conditions reflects *Hydra’s* delicate balance of reconciling an extremely active stem cell community with the maintenance of the - albeit simple - architecture of the organism (Bosch, et al. 2010). Histone ubiquitination, as another example, is a process well-known to be strongly conserved throughout evolution from yeast to mammals (Cao and Yan 2012; Wang, et al. 2017).

In stark contrast, the evolutionarily extremely short time frame of 44 years in the laboratory is characterized by significant adaptation of a number of biological processes, indicating that positive selection became the prevailing evolutionary force after this severe change of environmental conditions. Fitness adjustments to changes in laboratory conditions, is a phenomenon already observed in bacteria, worms and insects (Large, et al. 2016; Hoffmann and Ross 2018; Knöppel, et al. 2018). Our finding underlines *Hydra’s* extremely high adaptability to environmental changes providing the basis for its extraordinary sustainability over an evolutionary period of ∼200 million years (Schwentner and Bosch 2015).

Most adapted processes support the hypothesis that a significant proportion of selection in *Hydra* acts on the level of single stem cells rather than only at the level of individuals (Bosch, et al. 2010). Following a drastic environmental change, e.g., from the wild to the laboratory, cell clones may increase their prevalence in the cell population of an individual by fitness gains based on positive selection of: (i) “nucleic acid metabolic process” to improve nucleic acids availability for continued cell proliferation (Gross and Rotwein 2016; Zhu and Thompson 2019), (ii) “regulation of cellular response to growth factor stimulus” for faster cell cycle progression and higher division rates (Jones and Kazlauskas 2001; Fingar and Blenis 2004; Gross and Rotwein 2016), and (iii) faster “cell migration” (Fig.4A). The strongest sign for positive selection summarized by “generation of neurons”, however, points towards selection acting on the level of individuals as neuronal integration is essential for the fitness of a multicellular organism. Moreover, in *Hydra* cell migration to head and foot is associated with differentiation (Hager and David 1997; Siebert, et al. 2019).

Moreover, a remarkably high percentage of homozygous alternative SNV alleles was detected to have accumulated in *Hydra* cells in captivity. The majority of these homozygous variants found initially at the single-cell level could be confirmed at the whole-individual level, indicating that they are established with a significant MAF in the cell population.Two explanations appear to be plausible for this loss of heterozygosity: unidirectional “copy and paste” transfer of the mutated allele to its sister chromosome, e.g., as a consequence of the repair of a double strand break; or deletion of one non-mutated allele before or after the mutation event. Loss of heterozygosity was frequently observed in the context of continued mitotic cell divisions, such as many cancers as well as asexually unicellular organisms such as various yeast and funghi (Ryland, et al. 2015; James, et al. 2019; Dutta, et al. 2021). Our finding indicates that this phenomenon can also occur at a high frequency in a multicellular organism in the context of continued high and physiological mitotic activity. On the other hand, it may further underline the perspective on *Hydra* as an active stem-cell community sharing features of both multi- and unicellular organisms. It is tempting to speculate that this phenotype and the respective underlying molecular mechanisms have evolved *Hydra* specifically to accelerate adaptation to environmental changes.

Finally, we would like to emphasize that the presented genomic insights into the recent evolution of *H. magnipapillata* strain 105 primarily enhance our molecular understanding of *Hydra’s* extraordinary adaptability. Even though our data access only a narrow evolutionary time window, *a* mutation rate, comparable to that of human and mouse, generates sufficient genetic variability that allowed us to detect signs of negative selection over a longer time window before and of positive selection over only 44 years after a drastic environmental switch from wild to laboratory. Remarkably, the processes under strongest selection are closely related to those considered essential for the exapted, i.e. not brought about by natural selection, feature: *Hydra’s* non-senescence.

## Methods

For isolation of single cells, first head and foot were dissected. Dissociation of the remaining body column to single-cell suspensions and subsequent classification of cells was then conducted by an established protocol using a maceration solution containing glycerin, glacial acetic acid and water (David 1973). Sequencing was performed using Illumina’s next-generation sequencing methodology (Bentley, et al. 2008). In detail, 2 µg of amplified DNA was introduced into Illumina’s TruSeq DNA PCR-free library preparation. Paired end sequencing was conducted using Illumina HiSeq 2500 (high-throughput mode, 2x 101 cycles) and NovaSeq 6000 (Xp workflow, 2x 251 cycles) devices. Extraction of data was done using Illumina’s bcl2fastq v1.8.4 (HiSeq 2500) and v2.20.0.422 (NovaSeq 6000). The bioinformatics evaluation was done with the same workflow, programs and settings as a previous study that had determined somatic mutation rates in mouse and human (Milholland, et al. 2017). Briefly, Trim Galore was used for read trimming (https://github.com/FelixKrueger/TrimGalore), bwa mem for mapping (Li and Durbin 2009) against the *Hydra* reference genome (Chapman, et al. 2010), samtools for duplicate removal (Li, et al. 2009), GATK for indel realignment and base quality recalibration (Van der Auwera, et al. 2013), and VarScan2, MuTect and the UnifiedGenotyper for variant calling (intersection, required minimum read coverage of 20, Supplement Tables S1, S2, (Koboldt, et al. 2012; Cibulskis, et al. 2013; Van der Auwera, et al. 2013)).

For the estimation of the mutation rate per cell division, the number of mitoses since the zygote was assumed to be the sum of developing mitoses and continuous regeneration mitoses. For developing mitoses we used *log*_2_ (10^5^) = 17. For continuous regeneration mitoses, we used, given that I-cells and E-cells are known to divide every 24-30 h and 3-4 days, respectively (David and Campbell 1972; Campbell and David 1974), 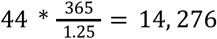 and 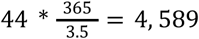, respectively.

For examining whole Hydra individuals, the Genomic DNA Isolation Kit from Norgen Biotek was used for DNA-extraction, the NEB Next Ultra II DNA Kit for library preparation (DNA input 10 ng) and an Illumina HiSeq 2500 device for paired end sequencing (2x 101 cycles). Extraction of data was done using Illumina’s bcl2fastq v2.20.0.422. Read trimming was conducted with Trimmomatic (Bolger, et al. 2014), read mapping with bwa mem (Li and Durbin 2009), removal of duplicates, indel realignment and base quality recalibration with GATK (Van der Auwera, et al. 2013), and clipped second mate overlaps using bamUtil (Jun, et al. 2015). Regarding the comparison of the buds vs their mother, for all buds, mapped reads were additionally downsampled to the lowest number of mapped reads among buds before variant calling. Variants were accepted if they were predicted by at least three of the following six callers: bcftools (Li 2011), freebayes (https://github.com/freebayes/freebayes), Mutect2 (Van der Auwera, et al. 2013), Platypus (Rimmer, et al. 2014), VarDict (Lai, et al. 2016) and VarScan2 (Koboldt, et al. 2012). For the comparisons of each individual vs reference, ‘germline’ variants were kept that were flagged accordingly. For the comparison of the buds vs their mother, only ‘somatic’ variants were kept that were flagged as somatic by Mutect2, VarScan2 and VarDict. For the remaining callers, bcftools, freebayes and Platypus, filters were adapted from the Bcbio-nextgen pipeline. Finally, only those somatic variants were kept for downstream analyses that either could be validated in the independent single-cell experiment analysis or had an FDR below 0.05 (Fisher-test). As a number of conducted tests for the multiple test correction, the number of respective germline variants (comparison vs reference) was used (Supplement Tables S1, S3). Our whole pipeline used for this workflow is available at https://github.com/Hoffmann-Lab/muvac. d_N_/d_S_ ratios were determined by the usual method (Nei and Gojobori 1986). As functional annotation for gene ontology analyses the Blast2GO (Conesa, et al. 2005) resource from the Hydra 2.0 Genome Project Portal was used (https://research.nhgri.nih.gov/hydra/download/functional_annotation/Blast2GO/blast2go_results.tsv).

## Data availability

Raw read data were deposited as ENA study with the accession PRJEB50344. The identified variants are available as browser accessible track hub at http://genome.ucsc.edu/cgi-bin/hgTracks?genome=Hm105_Dovetail_Assembly_1.0&hubUrl=https://genome.leibniz-fli.de/paras/hydra/hub.txt.

## Acknowledgements

We thank Ivonne Görlich, Cornelia Luge and especially Silke Förste for excellent technical assistance during this project. This work was supported by the Joachim Herz Foundation (Add-on Fellowships for Interdisciplinary Life Science, to A.S.).

## References

Bentley DR, Balasubramanian S, Swerdlow HP, Smith GP, Milton J, Brown CG, Hall KP, Evers DJ, Barnes CL, Bignell HR, et al. 2008. Accurate whole human genome sequencing using reversible terminator chemistry. Nature 456:53–59.

Boehm AM, Khalturin K, Anton-Erxleben F, Hemmrich G, Klostermeier UC, Lopez-Quintero JA, Oberg HH, Puchert M, Rosenstiel P, Wittlieb J, et al. 2012. FoxO is a critical regulator of stem cell maintenance in immortal Hydra. Proc Natl Acad Sci U S A 109:19697–19702.

Bolger AM, Lohse M, Usadel B. 2014. Trimmomatic: a flexible trimmer for Illumina sequence data. Bioinformatics 30:2114–2120.

Bosch TCG. 2009. Hydra and the evolution of stem cells. Bioessays 31:478–486.

Bosch TCG. 2007. Why polyps regenerate and we don’t: Towards a cellular and molecular framework for Hydra regeneration. Developmental Biology 303:421–433.

Bosch TCG, Anton-Erxleben F, Hemmrich G, Khalturin K. 2010. The Hydra polyp: Nothing but an active stem cell community. Development, Growth & Differentiation 52:15–25.

Busuttil RA, Rubio M, Dollé MET, Campisi J, Vijg J. 2006. Mutant frequencies and spectra depend on growth state and passage number in cells cultured from transgenic lacZ-plasmid reporter mice. DNA Repair 5:52–60.

Campbell RD, David CN. 1974. Cell Cycle Kinetics and Development of Hydra Attenuata: II. Interstitial Cells. Journal of Cell Science 16:349–358.

Cao J, Yan Q. 2012. Histone ubiquitination and deubiquitination in transcription, DNA damage response, and cancer. Frontiers in oncology 2:26–26.

Chapman JA, Kirkness EF, Simakov O, Hampson SE, Mitros T, Weinmaier T, Rattei T, Balasubramanian PG, Borman J, Busam D, et al. 2010. The dynamic genome of Hydra. Nature 464:592–596.

Cibulskis K, Lawrence MS, Carter SL, Sivachenko A, Jaffe D, Sougnez C, Gabriel S, Meyerson M, Lander ES, Getz G. 2013. Sensitive detection of somatic point mutations in impure and heterogeneous cancer samples. Nat Biotechnol 31:213–219.

Conesa A, Götz S, García-Gómez JM, Terol J, Talón M, Robles M. 2005. Blast2GO: a universal tool for annotation, visualization and analysis in functional genomics research. Bioinformatics 21:3674–3676.

Dańko MJ, Kozłowski J, Schaible R. 2015. Unraveling the non-senescence phenomenon in Hydra. Journal of Theoretical Biology 382:137–149.

David CN. 1973. A quantitative method for maceration of hydra tissue. Wilhelm Roux Arch Entwickl Mech Org 171:259–268.

David CN, Campbell RD. 1972. Cell Cycle Kinetics and Development of Hydra Attenuata: I. Epithelial Cells. Journal of Cell Science 11:557–568.

Domazet-Lošo T, Klimovich A, Anokhin B, Anton-Erxleben F, Hamm MJ, Lange C, Bosch TCG. 2014. Naturally occurring tumours in the basal metazoan Hydra. Nature Communications 5:4222.

Dong X, Zhang L, Milholland B, Lee M, Maslov AY, Wang T, Vijg J. 2017. Accurate identification of single-nucleotide variants in whole-genome-amplified single cells. Nature Methods 14:491–493.

Dutta A, Dutreux F, Schacherer J. 2021. Loss of heterozygosity results in rapid but variable genome homogenization across yeast genetic backgrounds. Elife 10:e70339.

Fingar DC, Blenis J. 2004. Target of rapamycin (TOR): an integrator of nutrient and growth factor signals and coordinator of cell growth and cell cycle progression. Oncogene 23:3151–3171.

Fujisawa T. 2003. Hydra regeneration and epitheliopeptides. Developmental Dynamics 226:182–189.

Gross SM, Rotwein P. 2016. Unraveling Growth Factor Signaling and Cell Cycle Progression in Individual Fibroblasts. The Journal of biological chemistry 291:14628–14638.

Hager G, David CN. 1997. Pattern of differentiated nerve cells in hydra is determined by precursor migration. Development 124:569–576.

Hemmrich G, Khalturin K, Boehm A-M, Puchert M, Anton-Erxleben F, Wittlieb J, Klostermeier UC, Rosenstiel P, Oberg H-H, Domazet-Lošo T, et al. 2012. Molecular Signatures of the Three Stem Cell Lineages in Hydra and the Emergence of Stem Cell Function at the Base of Multicellularity. Molecular Biology and Evolution 29:3267–3280.

Hoffmann AA, Ross PA. 2018. Rates and Patterns of Laboratory Adaptation in (Mostly) Insects. Journal of Economic Entomology 111:501–509.

Holstein TW, David CN. 1990. Cell cycle length, cell size, and proliferation rate in hydra stem cells. Developmental Biology 142(2):392–400.

Holtze S, Gorshkova E, Braude S, Cellerino A, Dammann P, Hildebrandt TB, Hoeflich A, Hoffmann S, Koch P, Terzibasi Tozzini E, et al. 2021. Alternative Animal Models of Aging Research. Frontiers in Molecular Biosciences 8.

James TY, Michelotti LA, Glasco AD, Clemons RA, Powers RA, James ES, Simmons DR, Bai F, Ge S. 2019. Adaptation by Loss of Heterozygosity in Saccharomyces cerevisiae Clones Under Divergent Selection. Genetics 213:665–683.

Jones OR, Scheuerlein A, Salguero-Gomez R, Camarda CG, Schaible R, Casper BB, Dahlgren JP, Ehrlen J, Garcia MB, Menges ES, et al. 2014. Diversity of ageing across the tree of life. Nature 505:169–173.

Jones SM, Kazlauskas A. 2001. Growth factor-dependent signaling and cell cycle progression. FEBS Letters 490:110–116.

Jun G, Wing MK, Abecasis GR, Kang HM. 2015. An efficient and scalable analysis framework for variant extraction and refinement from population scale DNA sequence data. Genome Research.

Kaliszewicz A, Lipińska A. 2013. Environmental condition related reproductive strategies and sex ratio in hydras. Acta Zoologica 94:177–183.

Klimovich A, Rehm A, Wittlieb J, Herbst E-M, Benavente R, Bosch TCG. 2018. Non-senescent Hydra tolerates severe disturbances in the nuclear lamina. Aging 10:951–972.

Knöppel A, Knopp M, Albrecht LM, Lundin E, Lustig U, Näsvall J, Andersson DI. 2018. Genetic Adaptation to Growth Under Laboratory Conditions in Escherichia coli and Salmonella enterica. Frontiers in Microbiology 9.

Koboldt DC, Zhang Q, Larson DE, Shen D, McLellan MD, Lin L, Miller CA, Mardis ER, Ding L, Wilson RK. 2012. VarScan 2: somatic mutation and copy number alteration discovery in cancer by exome sequencing. Genome Res 22:568–576.

Krishna S, Nair A, Cheedipudi S, Poduval D, Dhawan J, Palakodeti D, Ghanekar Y. 2013. Deep sequencing reveals unique small RNA repertoire that is regulated during head regeneration in Hydra magnipapillata. Nucleic Acids Research 41:599–616.

Lai Z, Markovets A, Ahdesmaki M, Chapman B, Hofmann O, McEwen R, Johnson J, Dougherty B, Barrett JC, Dry JR. 2016. VarDict: a novel and versatile variant caller for next-generation sequencing in cancer research. Nucleic Acids Research 44:e108–e108.

Large EE, Xu W, Zhao Y, Brady SC, Long L, Butcher RA, Andersen EC, McGrath PT. 2016. Selection on a Subunit of the NURF Chromatin Remodeler Modifies Life History Traits in a Domesticated Strain of Caenorhabditis elegans. PLOS Genetics 12:e1006219.

Li H. 2011. A statistical framework for SNP calling, mutation discovery, association mapping and population genetical parameter estimation from sequencing data. Bioinformatics (Oxford, England) 27:2987–2993.

Li H, Durbin R. 2009. Fast and accurate short read alignment with Burrows-Wheeler transform. Bioinformatics 25:1754–1760.

Li H, Handsaker B, Wysoker A, Fennell T, Ruan J, Homer N, Marth G, Abecasis G, Durbin R, Genome Project Data Processing S. 2009. The Sequence Alignment/Map format and SAMtools. Bioinformatics (Oxford, England) 25:2078–2079.

Lopez-Otin C, Blasco MA, Partridge L, Serrano M, Kroemer G. 2013. The hallmarks of aging. Cell 153:1194–1217.

Maroudas-Sacks Y, Garion L, Shani-Zerbib L, Livshits A, Braun E, Keren K. 2021. Topological defects in the nematic order of actin fibres as organization centres of Hydra morphogenesis. Nature Physics 17:251–259.

Martínez DE. 1998. Mortality patterns suggest lack of senescence in hydra. Exp Gerontol 33:217–225.

Martínez DE, Bridge D. 2012. Hydra, the everlasting embryo, confronts aging. Int J Dev Biol 56:479–487.

Milholland B, Dong X, Zhang L, Hao X, Suh Y, Vijg J. 2017. Differences between germline and somatic mutation rates in humans and mice. Nature Communications 8:15183.

Morley AA. 1995. The somatic mutation theory of ageing. Mutation Research/DNAging 338:19–23.

Müller WA. 1996. Pattern formation in the immortal Hydra. Trends in Genetics 12:91–96.

Nei M, Gojobori T. 1986. Simple methods for estimating the numbers of synonymous and nonsynonymous nucleotide substitutions. Mol Biol Evol 3:418–426.

Petersen HO, Hoger SK, Looso M, Lengfeld T, Kuhn A, Warnken U, Nishimiya-Fujisawa C, Schnolzer M, Kruger M, Ozbek S, et al. 2015. A Comprehensive Transcriptomic and Proteomic Analysis of Hydra Head Regeneration. Mol Biol Evol 32:1928–1947.

Rimmer A, Phan H, Mathieson I, Iqbal Z, Twigg SRF, Wilkie AOM, McVean G, Lunter G, Consortium WGS. 2014. Integrating mapping-, assembly-and haplotype-based approaches for calling variants in clinical sequencing applications. Nature Genetics 46:912–918.

Ryland GL, Doyle MA, Goode D, Boyle SE, Choong DY, Rowley SM, Li J, Bowtell DD, Tothill RW, Campbell IG, et al. 2015. Loss of heterozygosity: what is it good for? BMC medical genomics 8:45.

Sahm A, Bens M, Platzer M, Szafranski K. 2017. PosiGene: automated and easy-to-use pipeline for genome-wide detection of positively selected genes. Nucleic Acids Res 45:e100.

Schaible R, Sussman M, Kramer BH. 2014. Aging and potential for self-renewal: hydra living in the age of aging -a mini-review. Gerontology 60:548–556.

Schaible R, Scheuerlein A, Dańko MJ, Gampe J, Martínez DE, Vaupel JW. 2015. Constant mortality and fertility over age in <em>Hydra</em>. Proceedings of the National Academy of Sciences 112:15701–15706.

Schaible R, Ringelhan F, Kramer BH, Scheuerlein A. 2017. Hydra. In. Evolutionary and biological mechanisms for non-senescence: Cambridge University Press. p. 238–253.

Schwentner M, Bosch TC. 2015. Revisiting the age, evolutionary history and species level diversity of the genus Hydra (Cnidaria: Hydrozoa). Mol Phylogenet Evol 91:41–55.

Shimizu H, Sawada Y, Sugiyama T. 1993. Minimum tissue size required for hydra regeneration. Dev Biol 155:287–296.

Siebert S, Farrell JA, Cazet JF, Abeykoon Y, Primack AS, Schnitzler CE, Juliano CE. 2019. Stem cell differentiation trajectories in Hydra resolved at single-cell resolution. Science 365:eaav9314.

Stewart EJ, Madden R, Paul G, Taddei F. 2005. Aging and Death in an Organism That Reproduces by Morphologically Symmetric Division. PLOS Biology 3:e45.

Sturm A, Perczel A, Ivics Z, Vellai T. 2017. The Piwi-piRNA pathway: road to immortality. Aging Cell 16:906–911.

Sugiyama T, Fujisawa T. 1977. Genetic analysis of developmental mechanisms in hydra I. Sexual reproduction of Hydra magnipapillata and isolation of mutants. Development, Growth & Differentiation 19:187–200.

Supek F, Bosnjak M, Skunca N, Smuc T. 2011. REVIGO summarizes and visualizes long lists of gene ontology terms. PLoS One 6:e21800.

Tomczyk S, Fischer K, Austad S, Galliot B. 2015. Hydra, a powerful model for aging studies. Invertebr Reprod Dev 59:11–16.

Van der Auwera GA, Carneiro MO, Hartl C, Poplin R, Del Angel G, Levy-Moonshine A, Jordan T, Shakir K, Roazen D, Thibault J, et al. 2013. From FastQ data to high confidence variant calls: the Genome Analysis Toolkit best practices pipeline. Curr Protoc Bioinformatics 43:11 10 11–11 10 33.

Vijg J. 2021. From DNA damage to mutations: All roads lead to aging. Ageing Res Rev 68:101316.

Vogg MC, Beccari L, Iglesias Olle L, Rampon C, Vriz S, Perruchoud C, Wenger Y, Galliot B. 2019. An evolutionarily-conserved Wnt3/beta-catenin/Sp5 feedback loop restricts head organizer activity in Hydra. Nat Commun 10:312.

Vogg MC, Galliot B, Tsiairis CD. 2019. Model systems for regeneration: <em>Hydra</em>. Development 146:dev177212.

Wang P, Robert L, Pelletier J, Dang WL, Taddei F, Wright A, Jun S. 2010. Robust Growth of Escherichia coli. Current Biology 20:1099–1103

Wang L, Cao C, Wang F, Zhao J, Li W. 2017. H2B ubiquitination: Conserved molecular mechanism, diverse physiologic functions of the E3 ligase during meiosis. Nucleus (Austin, Tex.) 8:461–468.

Watanabe H, Schmidt HA, Kuhn A, Hoger SK, Kocagoz Y, Laumann-Lipp N, Ozbek S, Holstein TW. 2014. Nodal signalling determines biradial asymmetry in Hydra. Nature 515:112–115.

Wenger Y, Galliot B. 2013. RNAseq versus genome-predicted transcriptomes: a large population of novel transcripts identified in an Illumina-454 Hydra transcriptome. BMC Genomics 14:204.

Wolf AM. 2021. The tumor suppression theory of aging. Mechanisms of Ageing and Development 200:111583.

Yang Z. 2005. The power of phylogenetic comparison in revealing protein function. Proc Natl Acad Sci U S A 102:3179–3180.

Yousefzadeh M, Henpita C, Vyas R, Soto-Palma C, Robbins P, Niedernhofer L. 2021. DNA damage—how and why we age? Elife 10:e62852.

Zhu J, Thompson CB. 2019. Metabolic regulation of cell growth and proliferation. Nature reviews. Molecular cell biology 20:436–450

