## Supplement Text for "Non-aging despite high mutation rate – genomic insights into the evolution of *Hydra*"

### Supplement Text S1

#### Supplement Results

As described in more detail in the methods section of the main text, we used six different variant callers in 'somatic' to compare descendants (buds) against their mother. A variant was accepted if it was identified by at least 3 of these callers. The originally obtained variant numbers in the 6 older individuals (12.6-7.2 years; HM175_508, 734, 1170, 2003, 2419, 2450 ordered by age) examined varied between 27,200 and 28,171 (Supplement Table S3). Clearly less variants were found in the two youngest individuals examined (age 0.1 years; Hm_175bud1, bud3) with 24,499 and 24,501, respectively. This speaks against the hypothesis (discussed by Schaible, et al. 2015) that mutation-bearing cells would be preferentially ejected by the mother individual during budding. Furthermore, the two youngest individuals also showed the most similar variant profile (Fig. S1A). This suggests that budding is rather a random selection process of cells from Hydra' budding zone and resembles an evolutionary bottleneck leading to founder effects. Accordingly, the variant profile of the older individuals would have diverged further from that of the mother over time by genetic shift and selection. In general, we noticed unexpected overlaps between the variants identified in different individuals (Fig. S1B).

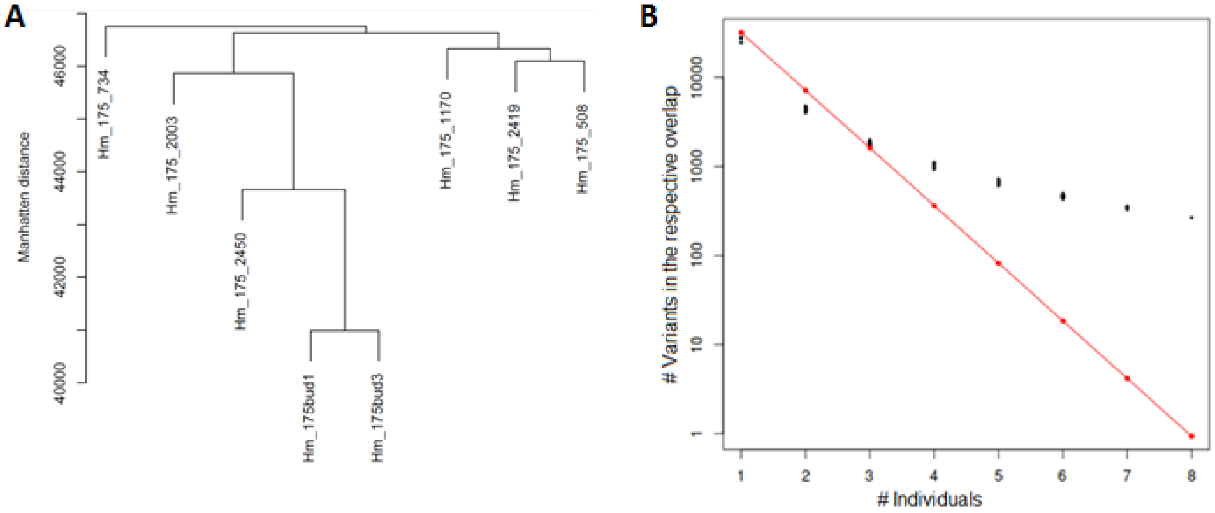


Fig. S1. (A) Clustering of the 8 descendants of the mother by their variant profile. The clustering was conducted based on Manhattan distances across all unique variant positions of the individuals. (B) Number of variants identified in 1 to 8 descendants of the mother. For each number of individuals displayed on the x-axis all sets of individuals of the respective size were formed and the number of SNVs shared by all members of the set was determined. Each black point represents the number of shared variants of one of those sets. The red points represent the expected number of shared variants according to a simple stochastic model where we assumed a total number of variant positions $K$ in the mother (140,855), a probability $P$ that the respective variant allele is transmitted during a budding event (fraction of mother cell transmitted to the bud: 0.225) and the resulting number $E = P^n * K$ representing the variants to be expected in the overlap of $n$ individuals.

The simplest - most parsimonious - explanation of this phenomenon would be that those variants would have existed already in the mother with a too low minor allele frequency (MAF) to be detected with the available read coverage (sequencing depth) and the applied filter criteria (see Methods). For confirmation we inspected the sequencing reads from the mother at the 267 variant positions shared by all 8 buds. We often found 1-2 reads representing the alternative alleles in support of our assumption.

Next, we asked whether the higher than random overlap of variants between buds (Fig. S1) could be the result of positive selection acting on the buds. Following this idea cells carrying a shared allele would have a selective advantage during or after budding leading to expansion of this cell clone and respectively the MAF of the variant in the bud. In this case it could be expected that the ratio of nonsynonymous to synonymous variants should be the greater the more buds share it. However, we found the opposite. While the number of and synonymous variants indeed displayed significantly different trajectories depending on the number of individuals sharing them ($p<2.2*10^{-16}$, Fig. S2) the ratio shifted in favor of synonymous variants. This means that variants shared by more buds would be subject to stronger negative selection, contrary to our assumption.

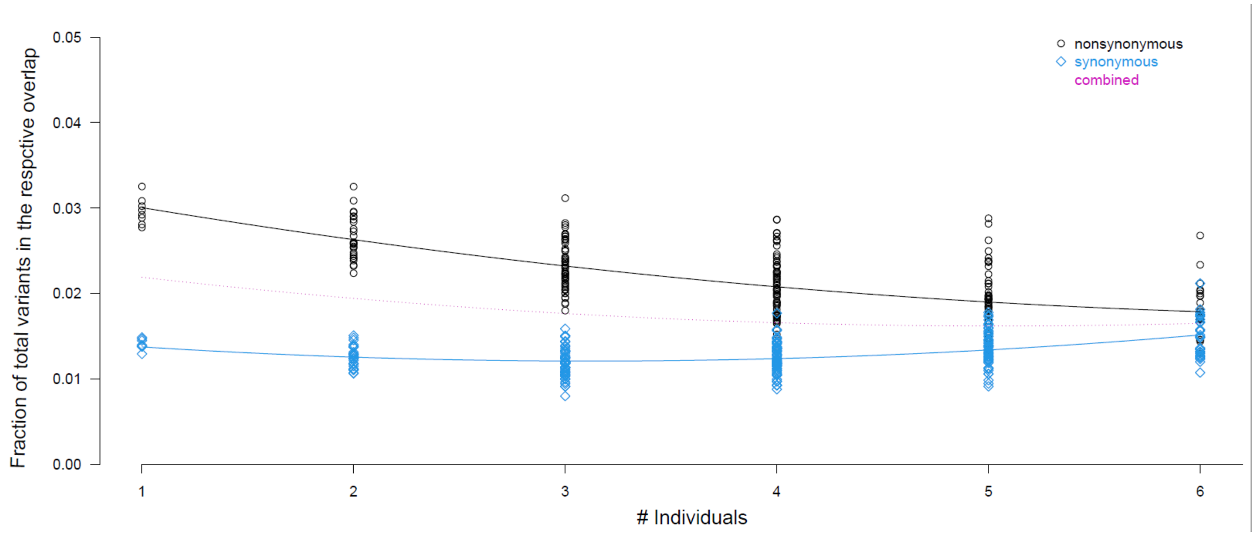


Fig. S2. Nonsynonymous and synonymous variant fractions depending on the number of individuals sharing them. Each dot represents the number of SNVs shared by all members of a set of individuals of the respective size. Variants in coding sequences shared by 7 or 8 individuals were not included because their numbers were with 10 to 14 too low to meaningfully determine proportions.

In search of another explanation of nonrandom variant sharing, we again took a closer look at samples of the 267 variants shared by all 8 buds. Now inspecting the sequencing reads from the buds, we got the impression that most of the variants had a MAF of about 0.5. In principle, a MAF of about 0.5 is an indicator of heterozygosity. This interpretation is supported by the fact that many variants were clustered in close vicinity and formed physical haplotypes, i. e. were found on the same sequencing read (Fig S3).

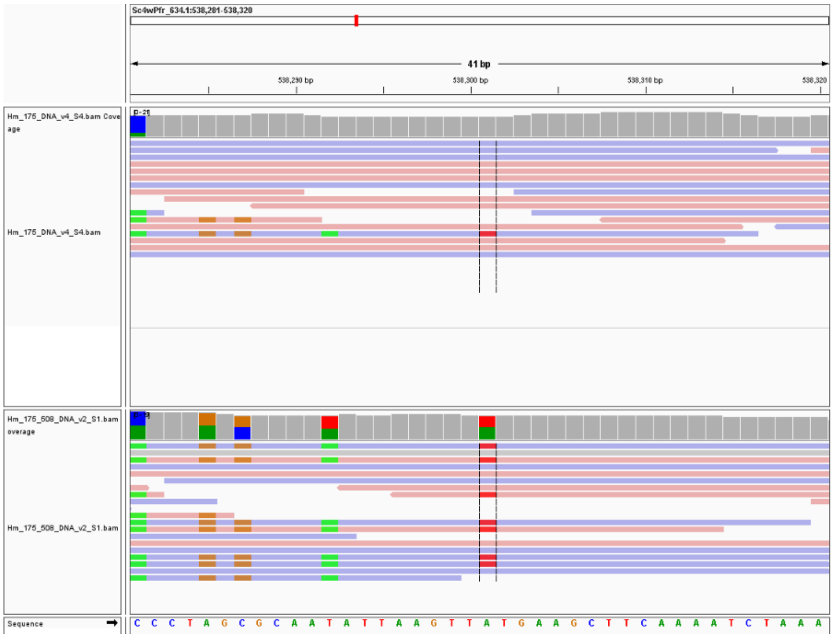


Fig. S3. Example of a variant (red, center) that was shared by all 8 buds in the comparison against their mother. Displayed is an igv genome browser image showing the read coverage of the mother (top) in this selected genome region, the read coverage of one of the buds at this position (middle) and the *Hydra* reference genome sequence (bottom). Variants are highlighted by the respective colors of the substituting bases (c.f. with color code of the reference sequence). The red colored variant (A T) was not only displayed by the bud shown here but by all 8 buds.

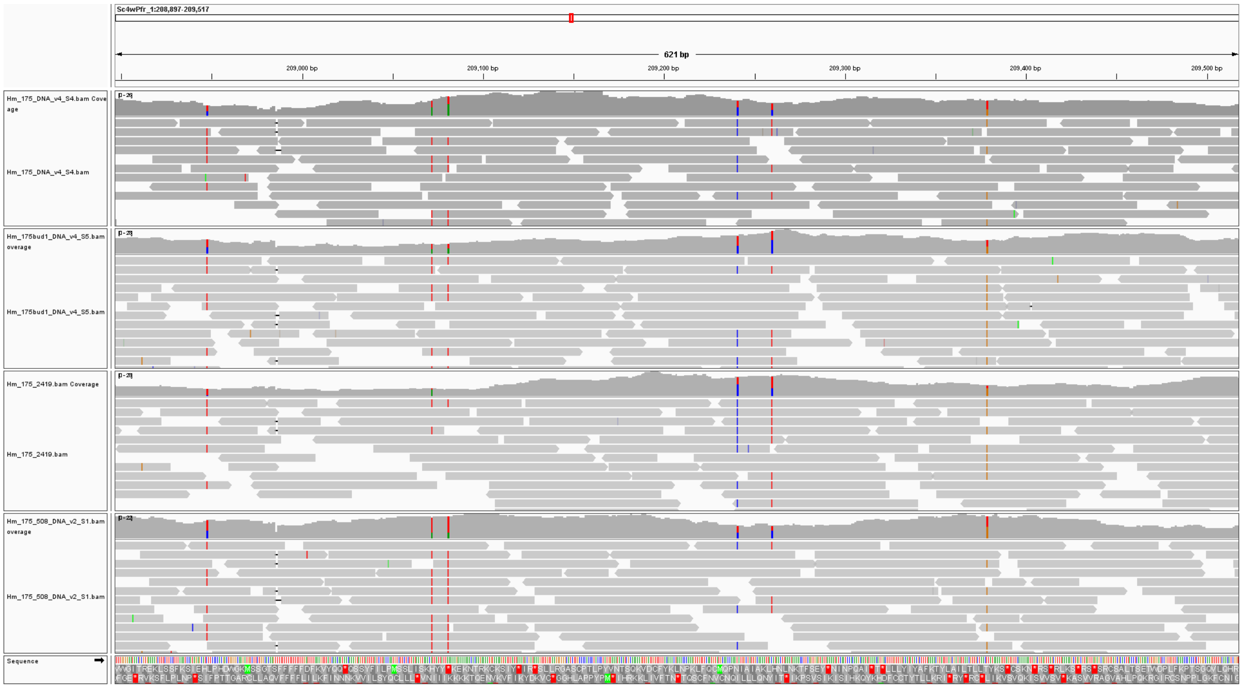


Fig. S4. Example of a variant identified against the reference genome of *Hydra*. Displayed is an igv genome browser image of a selected genome region showing the read coverage of the mother at the top, and that of three of her buds below. Colored lines indicate deviations from the reference genome, whose sequence is shown at the bottom.

Sensitized by these insights, we also examined samples of variants that were called in the comparison against the reference genome variants. Again, often long stretches of coupled variants - physical haplotypes - showed up, mostly with a MAF of about 0.5 (Fig. S4). Of note, in this comparison, a 2-3 orders of magnitude greater number of variants was found (Ø 4.5m, Supplemental Table S3) than in the comparison of the individuals against their mother (Ø 27k) and that of the single cells (Ø 14k and 3k, for I- and E-cells, respectively). These large differences can be explained only to a small extent by the different genome proportions that were examined in these comparisons due to the varying coverage and filtering requirements: Ø 77.6% in the comparison against the *Hydra* genome reference;Ø 51.9% in the comparison of the buds against the mother; ,Ø 9.6% in the I-cell comparison; Ø 6.9% in the E-cell comparison (Supplement Tables S2 and S3). The much larger number of variants found in the comparison against the monoallelic reference genome suggests that the vast majority of them were accumulated over a substantially longer evolutionary distance, i.e., not just the few decades of captivity of the *Hydra* strain 105 that the other two comparisons represent. Instead, in agreement with the observations described above, it is reasonable to assume that the vast majority of them arose during the preceding period in the wild and were heterozygous from the zygote onward.

Based on these observations, we formulated a new hypothesis to explain the high overlap of variants found in the comparison of the buds against their mother. Thus, most overlapping variants would be mere extreme cases of the sampling distribution, of the total of several million multiallelic positions in the mother. Consequently, the majority of those overlapping variants would also be in fact heterozygous germline variants with a true MAF of 0.5. If we assume this to be the case then we assess the measured MAF in the mother as underestimated and the MAFs in the buds could be seen as estimators of the true MAF of the mother. Consequently the underestimation in the mother would be more secure if more buds would deviate from the mother. So the MAF observed in the buds should increase with the number of individuals sharing a variant. In agreement with this, the mean MAF shifts from 0.22 in variants found in only 1 individual to 0.46 in variants found in 8 individuals (Fig. S5), suggesting that most the latter are indeed likely to be heterozygous germline variants of the entire strain. Again in line with the observations mentioned above, we also found a bimodal MAF distribution for the mother where one peak stands for zero reads and one peak for one or two reads representing the alternative allele. The ratio of the peaks gradually shifts depending on how many individuals share the variant: for variants that were found in only one individual, the mother most often shows no reads supporting the alternative allele, for variants shared by 8 individuals, however, one or two mother reads supporting the alternative allele in the majority of cases (Fig. S5). This is what we expect following the assumption that the true MAF in the mother equals that of the buds, because less extreme underestimation, within the boundaries still accepted by the callers, has to occur at a higher frequency than more extreme underestimations with 0 reads.

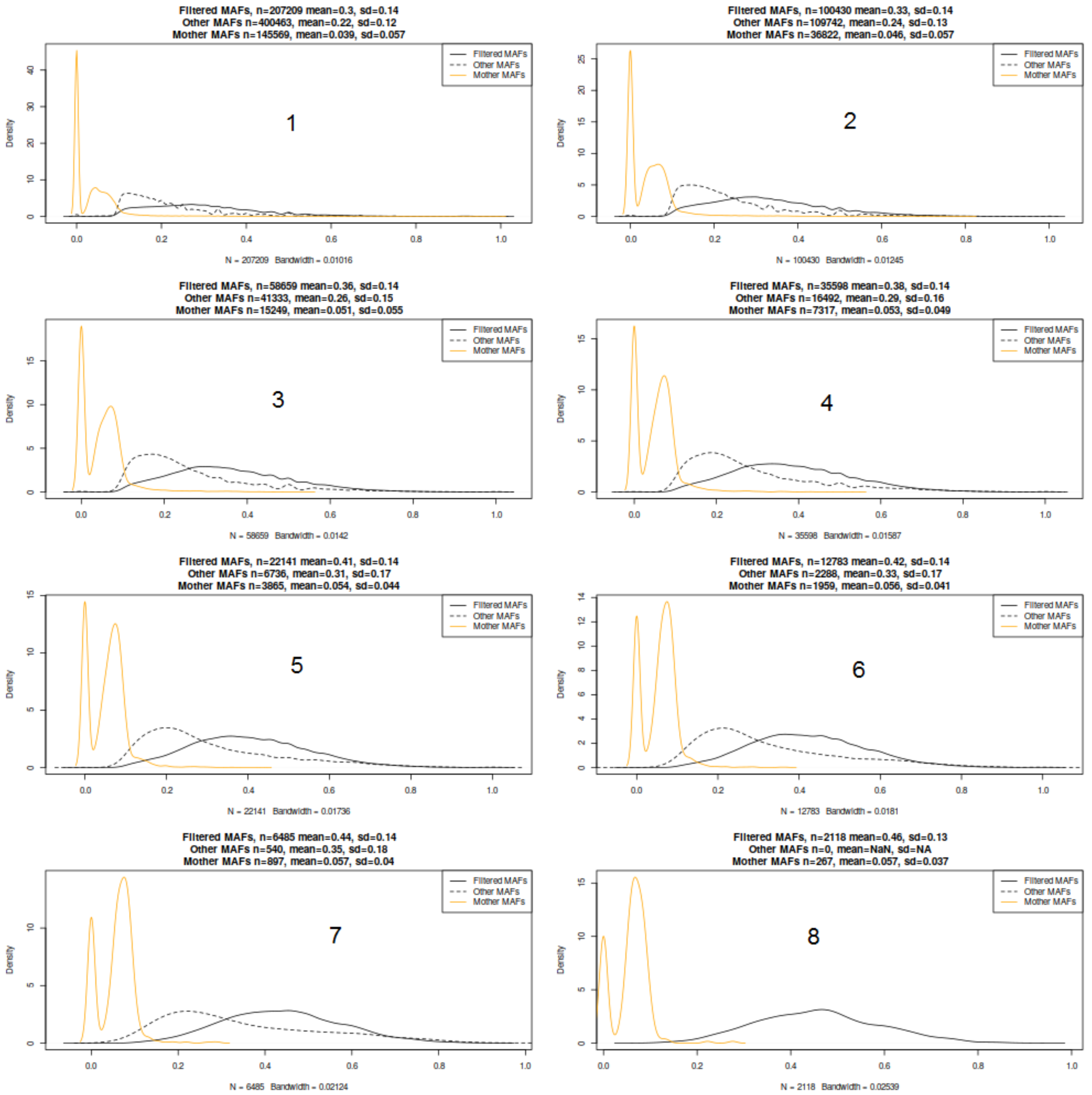


Fig. S5. Minor allele frequency (MAF) distributions for variants shared by 1 to 8 buds in the comparison against their mother. ‘Filtered MAFs’ stand for variants that were identified by at least three of the six variant callers used according to our methodical approach. ‘Other MAFs’ stand variants that were identified by only one or two callers.

If we accept that wrong MAF estimation due to read sampling during sequencing can have a significant influence, we have to conclude that the underestimation of MAF in the mother is just one special case. Conversely, overestimation in buds is also possible. We would expect that overestimation and underestimation of MAF due to sampling errors are in principle more likely to happen the lower the read coverage at the position in question is. Consequently we expect to see the highest MAF differences between buds and mother in regions sparsely covered, and the higher the coverage in a variant, the smaller the MAF differences. What we do not expect under the assumption that MAF differences are due to sampling errors are highly covered genome regions with high MAF differences. These expectations are fulfilled to a very large extent when examining the MAF differences as a function of read coverage (Fig. S6), suggesting that most of the supposed variants are in fact stochastic artifacts.

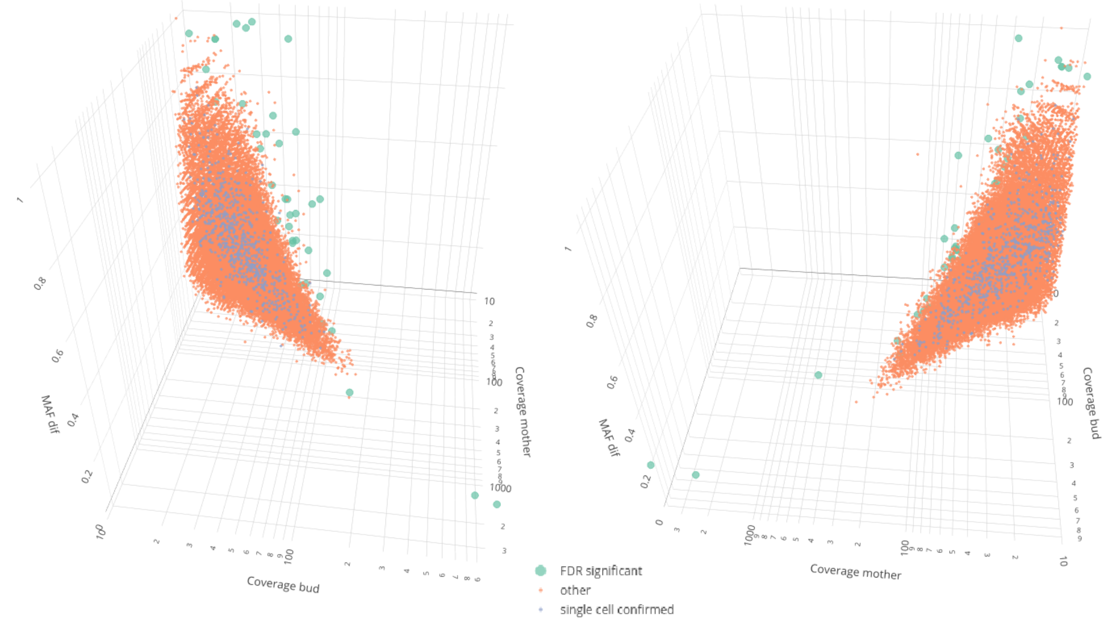


Fig. S6. Variants identified in the individuals in the comparison of the individuals against their mother. Display is the absolute difference in the minor allele frequency (MAF) between the bud and the mother in dependence of the read coverage of mother and bud.

The most likely determining factor in the occurrence of these stochastic artifacts is the very high number of multiallelic positions throughout the strain. By far the most common case is likely to be heterozygous positions arising from the zygote. However, over- or underestimation of the MAF can basically occur at all positions with more than one allele, e.g., also cell clones with specific variants that stand for a larger part of the cell population of the strain. We can estimate the number of the multiallelic positions by the number of identified variants in the comparison against the reference genome. If we reanalyze the variants identified in comparison to the mother on the basis of a simple urn model with Fisher's test we come to a median P-value of the variants of 0.04. If we take this as the average alpha error and our estimate of the heterozygous variants in the strain (ca. 3.4-5.2 million depending on the individual), the number of variants expected just by multiple testing (ca. 130,000 – 200,000) is even significantly higher than the number of variants actually identified (ca. 24,500-28,000). The reason for the latter is presumably that we required that multiple variant callers with sometimes very different methodological sets had to confirm a variant. In any case, it can be assumed that the majority of variants identified compared to the parent represent expectable stochastic noise due to multiple testing. For this reason, we decided to list only those variants in the main section that showed significant differences also after multiple test correction of the p-value resulting from Fisher's test (n=46, green points in Fig. S6) or were independently confirmed by the single cell experiment (n=746, blue points in Fig. S6, Supplement Table S3). These 809 variants in total found for the comparison of the individuals against their mother correspond to 505 unique variant positions.

In conclusion, these initial data-mining efforts have provided us with tool to reliably detect variants originated either before or after the last sexual reproduction leading to the asexual strain 105 of *H. magnipapillata* and encouraged us to concentrate on the identification of evolutionary forces and targets shaping its biology.

#### Supplement Methods

Curve fitting for Fig. S2 was performed in R version 4.1.2 (http://www. R-project.org) by minimizing the least squares deviation of a set of regression functions to the data. The best fitting function was determined by the Akaike information criterion. For comparison of two groups, the sum of the residual sum of squares of the individual regressions is then compared to that of the regression for the combined data using the F-test (Motulsky and Ransnas, 1987).

Motulsky HJ, Ransnas LA (1987) Fitting curves to data using nonlinear regression: a practical and nonmathematical review. FASEB J 1: 365–374
